# The influence of force on the encoding and perception of affective touch

**DOI:** 10.64898/2026.01.15.699730

**Authors:** Sophia Faresse, Luidgi Thrace, S. Hasan Ali, Adarsh Makdani, Francis McGlone, Andrew G. Marshall, Paula D. Trotter, Rochelle Ackerley

## Abstract

Stroking touch has been shown to be most pleasant at intermediate velocities of 1-10 cm.s^−1^, which relates well to the activity of C-low threshold mechanoreceptors (C-LTMRs), also called C-tactile afferents in humans, that have been implicated in encoding positive affective touch. This well-established finding has been demonstrated at gentle stroking forces (typically 0.4 N peak normal force), yet few studies have investigated the effect of force on the perception of stroking touch or on the activity of C-LTMRs. Presently, we investigated the pleasantness and intensity of stroking touch (0.3, 1, 3, 10, 30 cm.s^−1^) at different forces (0.2, 0.4, 0.8, 1.2, 1.6, 2.0 N) on hairy forearm skin and compared this to responses from C-LTMRs during stroking touch at the same velocities, but over fewer forces (0.05, 0.4, 1.5 N). We found significant effects of stroking velocity and force for both tactile pleasantness and intensity ratings. Pleasantness showed the typical inverted-U shaped relationship over stroking velocities, but this was modified by force: higher forces significantly decreased pleasantness at faster velocities. Conversely, tactile intensity increased linearly with both increasing velocity and force. Recordings from seven C-LTMRs showed that their firing frequency changed with stroking velocity, but that they were strongly modulated by force. Overall, we demonstrate the profound effect that force has on the perception of stroking touch, where tactile pleasantness and intensity appear to be a multi-faceted construct that are related to C-LTMR and Aβ-LTMR activity, respectively, but that the firing in individual mechanoreceptor populations cannot fully account for percepts.

**Summary:**

1. The skin is highly sensitive to small mechanical deformations, including dynamic touch and different forces.
2. We obtained perceptual ratings for touch pleasantness and intensity over different stroking velocities and forces applied to the forearm. This was also compared to microneurography recordings from C-low threshold mechanoreceptors (C-LTMRs).
3. Building on previous work, we find that it is not only stroking velocity that affects pleasantness and intensity perception, but also stroking force. Pleasantness ratings decreased more at higher forces when the stroking was faster. C-LTMR firing showed that force differences were readily encoded, although the same force-velocity interaction was not seen.
4. This work expands the literature on affective touch to encompass the effects of force, which clearly modulates pleasantness.
5. C-LTMR force and velocity information likely combines with information from all types of mechanoreceptors, including those from the fascia, to produce the resulting percept.

Abstract figure.
By combining perceptual experiments with neuronal recordings, we obtained ratings of perceived pleasantness and intensity during stroking touch applied at different velocities and forces, and recorded the single-unit activity from 7 C-low threshold mechanoreceptors during similar tactile stimulation. We found that the perception of pleasantness and intensity was significantly influenced not only by the velocity of the stroking, but also by the force applied. Pleasantness ratings decreased more at higher forces when the stroking was faster. C-LTMR activity showed that differences in force were readily encoded, although the same force-velocity interaction was not observed. Our results support the hypothesis that the perception of the pleasant aspect of touch may be influenced by the activity of C-LTMR afferents, but arises from the combined activity from other low threshold fast-conducting mechanoreceptors.

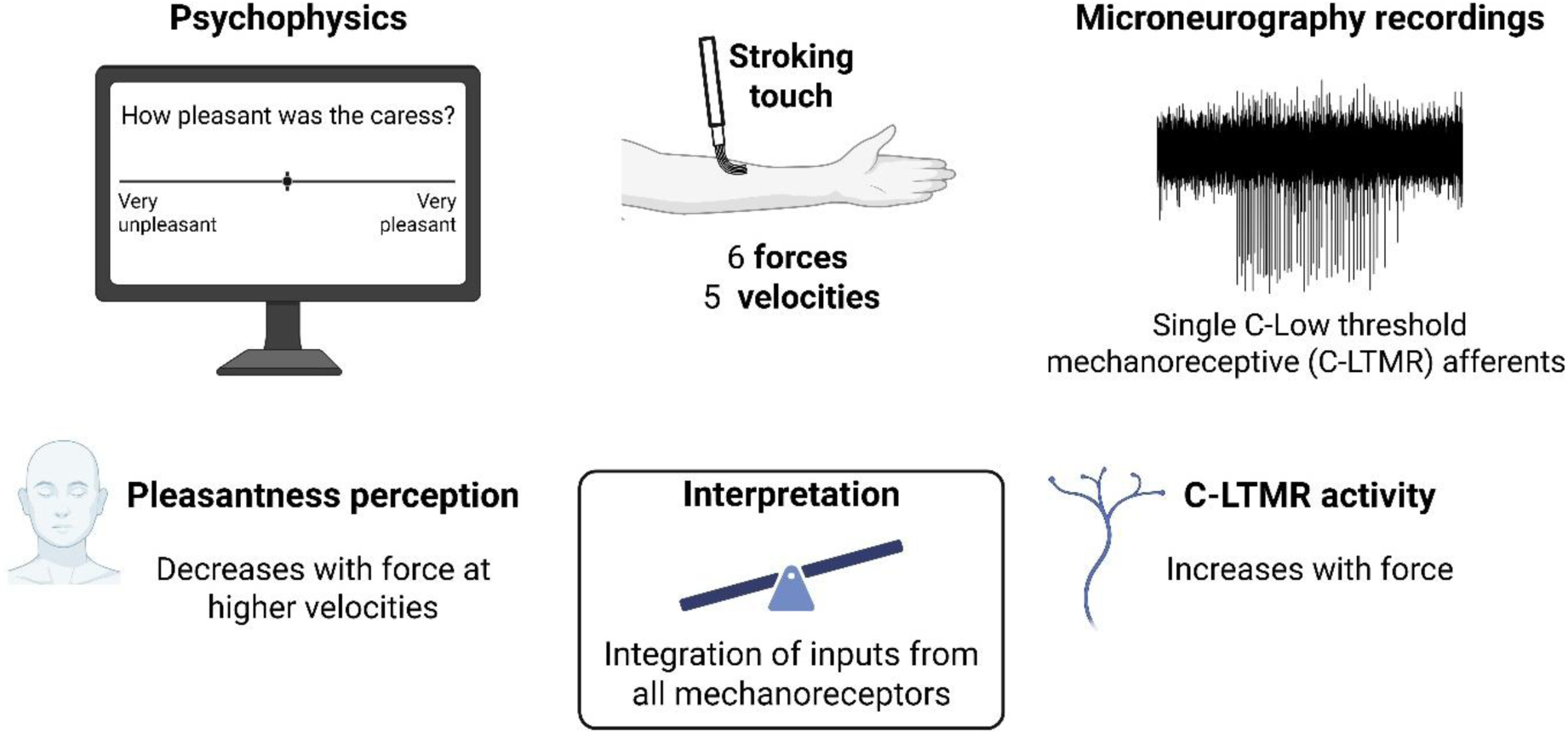

## Introduction

Touch encoding begins in the skin, where low-threshold mechanoreceptors (LTMRs) located at the ends of afferent nerve fibres transmit specific components of mechanical signals to the central nervous system. Tactile perception is based upon this input, combined with the contextual, cognitive and emotional state. LTMRs are all exquisitely sensitive to skin deformations and vary in type and density depending on the area of the body (Vallbo & Johansson, 1984; Vallbo *et al*., 1995; Corniani & Saal, 2020). While the discriminative aspects of touch (e.g. knowing where, when, for how long) are encoded by fast-conducting Aβ-LTMR fibres, the positive affective (emotional) aspects of touch are believed to be reinforced by C-tactile (CT) afferents in humans, also known more generally as C-low threshold mechanoreceptors (C-LTMRs) (McGlone *et al*., 2014; Schirmer, Croy *et al*., 2023). LTMRs all respond to low-force, dynamic touch, where microneurography studies in humans have shown how Aβ-LTMRs encode various factors, such as indentation force and velocity (Knibestöl, 1973, 1975), stroking velocity (Essick & Edin, 1995; Edin *et al*., 1995), and dynamic force directionality (Birznieks *et al*., 2001), mainly in the glabrous, non-hairy skin of the hand. However, less is known about how forces are encoded and perceived in hairy skin, which is important to consider with respect to its role in receiving affective touch (McGlone *et al*., 2014; Schirmer, Croy *et al*., 2023).

Hairy skin has been investigated extensively on the forearm, in many different studies that have applied stroking stimuli at various speeds, focussing on the relationship between velocity and the pleasantness of touch. Initial work found that there was an inverted-U shaped relationship between stroking velocity and tactile pleasantness, where very slow and very fast speeds are rated as less pleasant (Essick *et al*., 1999). This was followed-up by a microneurography study recording from C-LTMRs innervating the forearm that linked their firing with the perceived pleasantness of stroking touch (Löken *et al*., 2009). Here, the authors showed the tactile pleasantness of stroking followed an inverted-U shaped curve, where velocities between 1-10 cm.s^−1^ were perceived as the most pleasant, which was shown to relate to the mean firing frequency of C-LTMRs. Further, the authors also found a monotonic increase in the firing frequency of various Aβ-LTMR fibres from 0.3-30 cm.s^−1^, reflecting that they linearly encode increasing stroking velocity. A similar study found the same results with regards to the relationships in C-LTMR firing at neutral temperature (akin to skin temperature) and also for Aβ-LTMR hair follicle afferents (Ackerley, Wasling *et al*., 2014). A more recent study also found this inverted-U shaped relationship for C-LTMRs on the hairy skin of the leg (Löken *et al*., 2022). Two other investigations demonstrated the high sensitivity of Aβ-LTMRs to stroking, where there was again an exponential increase in the firing of fast-adapting (FA) and slowly-adapting (SA) Aβ-LTMRs (Essick & Edin, 1995; Edin *et al*., 1995).

The relationship between the perception of tactile pleasantness and microneurography recordings has led to the affective touch hypothesis (Vallbo *et al*., 2009; Olausson *et al*., 2010) that postulates a dominant role for C-LTMRs in pleasant touch perception, which was furthered by the social touch hypothesis (Olausson *et al*., 2010; Morrison *et al*., 2010) that specifically implicated C-LTMRs in reinforcing human-to-human interactions. Many papers followed that have shown a clear relationship between stroking velocity and perceived pleasantness, where the intermediate speeds are more pleasant (e.g. Ackerley, Carlsson *et al*., 2014; Croy *et al*., 2021; Cruciani *et al*., 2021). However, there has been little consideration of how force influences C-LTMR firing and affective touch perception.

All the above microneurography studies investigating LTMR firing and stroking velocity have used a type of rotary tactile stimulator (RTS; Essick *et al*., 1999, 2010). This is typically calibrated to apply a maximum normal force of 0.4 N during stroking over the skin and many psychophysical studies have also used this stroking force. Löken *et al*. (2009), tested two normal forces, namely 0.2 and 0.4 N, and no significant differences were found between these forces for the C-LTMR afferents, nor in the pleasantness ratings. Conversely, Aβ-LTMR firing frequencies have been found to increase with force for some SA and FA Aβ-LTMRs afferents, although the forces applied did not exceed 0.4 N (Essick & Edin, 1995; Edin *et al*., 1995; Löken *et al*., 2009). In terms of tactile perception, differences have been found in pleasantness with increasing force between textures, where soft-smooth textures (e.g. a brush) tend to have a constant level of pleasantness over different forces, whereas rougher-harder textures tend to decrease in pleasantness with increasing force (Cascio *et al*., 2008); however, the forces were not well controlled, as stroking was manually-applied at light, medium, or high forces. A further study using the RTS found that pleasantness generally decreased with stroking force, but this was less clear for soft-smooth textures and the maximum force tested was 0.9 N (Essick *et al*., 2010). A more recent study explored the effect of more intense forces on stroking touch, testing the typical range of stroking velocities (0.3-30 cm.s^−1^) at RTS-set forces of 0.05, 0.4, and 1.5 N (Ali *et al*., 2023). They found that for all velocities, touch was perceived as more pleasant at 0.4N compared to 0.05 and 1.5 N, and 0.05 N was significantly more pleasant than 1.5 N. Taking all these into consideration, we were thus interested in testing a wider range of forces.

Touch is multi-faceted and participants can easily rate various percepts, other than pleasantness (Jönsson *et al*., 2015; Sailer *et al*., 2020; Schirmer, Cham *et al*., 2023). One other relatively common percept to measure in skin stroking studies is tactile intensity, which has been generally found to increase linearly over the typical range of velocities tested (Triscoli *et al*., 2013; Jönsson *et al*., 2015; Sehlstedt *et al*., 2016; Schirmer, Cham *et al*., 2023). Intensity appears to relate more to the firing of Aβ-LTMRs, which increase their activity monotonically with stroking velocity.

We investigated how stroking force affects pleasantness and intensity perception over different stroking velocities (Experiment 1), as well as relating this to the firing of C-LTMRs in a similar paradigm (Experiment 2), as C-LTMR activity has been directly linked with pleasantness and little is known about their force encoding. We hypothesised that perception would vary with both the force and the velocity, but that this would be in different ways for pleasantness and intensity. Further, we hypothesised that C-LTMR activity would also be modulated directly by stroking velocity and force, where the typical inverted-U shape would be seen for velocity, but that firing frequency would increase monotonically with force, as has been shown for C-LTMR firing during short normal indentations with calibrated monofilaments (Middleton *et al*., 2022), and during stroking for other Aβ-LTMRs (Knibestöl, 1973, 1975; Essick & Edin, 1995; Edin *et al*., 1995; Löken *et al*., 2009).

## Materials and Methods

### Experiment 1: Psychophysical study into the perception of stroking at different velocities and forces

#### Ethical approval and participants

The experiment was approved by the ethical committee Comité de Protection des Personnes Île-de-France X (no. 2023-A01990-45) and was conducted in accordance with the guidelines set out in the Declaration of Helsinki (1964) and its later amendments, apart from pre-registration in a database. We recruited 21 adult participants (aged between 19 and 29 years old, 6 males) who participated in the experiment at the Centre for Research in Psychology and Neuroscience in Marseille (UMR 7077). Participants were recruited by mailing list and word of mouth. Exclusions criteria included pregnancy, being placed under legal protection, inability to understand the information and consent, and having a self-reported disorder that could alter tactile sensitivity (e.g. diabetic neuropathy, skin condition, psychiatric or neurological disorders). Participants received verbal and written information prior to agreeing to participate and they all gave written informed consent. They received a remuneration of 10 euros for their time.

#### Stroking stimuli

We used a rotary tactile stimulator (RTS; Dancer Design, UK) robot to apply stroking stimuli to the middle of the forearm (determined with a measuring tape between the wrist fold and cubital fossa), between the ventral and dorsal aspects, as aligned with the thumb (Fig. 1A) at controlled velocities and calibrated normal forces. We used a flat soft artists’ brush that had a width of 3 cm for the hairs, with a density of ∼1 cm. We applied five stroking velocities: 0.3, 1, 3, 10, and 30 cm.s^−1^, over six forces: 0.2, 0.4, 0.8, 1.2, 1.6, and 2 N. The stroking was always applied in a proximal-to-distal direction. The normal forces were calculated from a dynamic stroking force curve that was calibrated up to the maximum force. Force curves were saved for each stroking trial. Stroking velocities were applied in a pseudorandomised order and each velocity-force combination was repeated three times. This gave a total of 90 applied stimuli and the experiment was completed in 1 hour.

**Figure 1:**
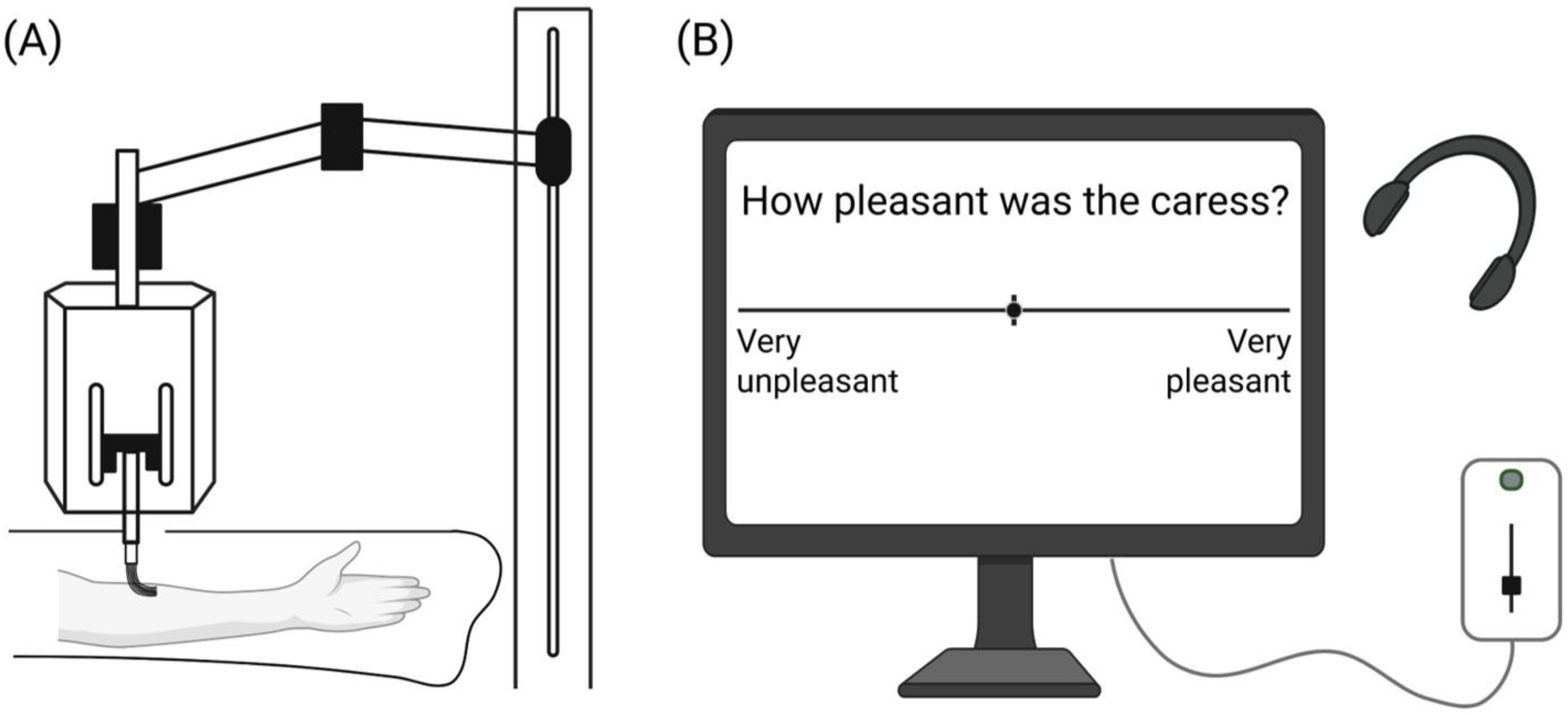
Diagram of Experiment 1 setup. (A) Diagram of the rotary tactile stimulator robot. Stroking touch was applied to the middle of the left forearm. (B) Diagram of the slider device and the screen used to collect participants’ ratings by moving the cursor displayed on the screen. A rating of pleasantness is shown as an example; the same design was used for intensity ratings (“How intense was the caress?” from “Very weak” to “Very strong”).

#### Procedure

Participants were seated comfortably in a chair, in front of a computer screen. Their left forearm was secured using a vacuum cushion to prevent movement and keep the forearm in a fixed and comfortable position. In their right hand, they held a visual analogue scale (VAS) slider device that was used to move a cursor on the screen to give tactile pleasantness and intensity ratings. A curtain was used to prevent the participant from seeing their forearm and noise-cancelling headphones (Bose) were used to limit external sounds. The setup is presented in Figure 1. For each participant, the RTS was calibrated to apply the chosen forces before the experiment started. For the experiments, the participant was instructed to pay attention to the moving touch and then rate its pleasantness and intensity after each stroke, using two separate VASs displayed on a computer screen. After a stroking stimulus had terminated, the first VAS to rate tactile pleasantness appeared (‘How pleasant was the caress?’), where the touch was rated from ‘Very unpleasant’ (corresponding to −10) to ‘Very pleasant’ (corresponding to +10) (Fig. 1). After they chose their response and clicked a button to enter it, a second scale appeared where they rated the intensity of the touch (‘How intense was the caress?’) from ‘Very weak’ (corresponding to −10) to ‘Very strong’ (corresponding to +10). Once the second rating had been completed, the next stimulus was delivered.

#### Analysis

Preprocessing and visualizations of the data were conducted with Python (Jupyter Notebook, version 3.11.4 packaged by Anaconda). The statistical analyses were conducted in RStudio (R v4.4.0), where linear and generalized linear mixed model analyses, as well as regressions, were conducted with the lme4, lmerTest, glmmTMB, and post-hoc comparisons with the emmeans packages. As per previous literature, the decimal logarithm (log10) of the stroking velocity was used (Löken *et al*., 2009), to make the spacing between velocities equal. Alpha error was fixed at p < 0.05. Normal distribution of the residuals of the linear mixed models were assessed visually using histograms and QQ plots. Effect sizes were measured using partial eta squared (η²ₚ). Ratings were analysed using separate linear mixed-effects models. The dependent variable was the pleasantness or intensity rating. Force, velocity, and their interaction were included as fixed effects. The model included a random intercept for participant and a random slope for velocity across participants. Bonferroni-corrected post-hoc tests were conducted to compare between stroking velocities and forces, and further contrast comparisons were used to investigate interaction effects, namely between ratings of forces at each stroking velocity. A linear mixed model was also used in the analysis of the force recorded by the RTS. The dependent variable was the recorded force. Set force, velocity, and their interaction were included as fixed effects. The model included a random intercept for participant. Bonferroni-corrected post-hoc tests were conducted to compare between stroking velocity and set force and for interactions, contrast comparisons were conducted for each force, namely between the velocity of 30 cm.s^−1^ (fastest), as compared to all the other velocities.

### Experiment 2: Microneurography recordings from C-LTMR afferents to stroking at different velocities and forces

#### Ethical approval and participants

Experiment 2 was approved by the local ethical committee of Liverpool John Moores University (no. LJMU REC 14/NSP/039) in the UK and by the Comité de Protection des Personnes Sud-Ouest et Autre Mer III (no. 2021-A01604-37) in France. The research was conducted in accordance with the guidelines set out in the Declaration of Helsinki (1964) and its later amendments, apart from pre-registration in a database for the UK. As typically found in microneurography studies studying single unit cutaneous recordings from forearm hairy skin, the success rate is rather low and the data gained can be sparse. The data presented in the current study includes recordings from four healthy adult participants (2 females (20 and 39 years) and 3 males (20, 26, and 26 years)), where it was possible to run the full protocol on seven C-LTMRs. All participants received verbal and written information, and signed an informed consent form.

#### Stroking stimuli

The set-up was a duplicate of that in Experiment 1, where an RTS stroked over the forearm skin, depending on where the receptive field of a C-LTMR was found. The middle of the stroke was centred over the hotspot of the receptive field. The same five stroking velocities were used (0.3, 1, 3, 10, 30 cm.s^−1^), although the forces tested were a little different, namely a very low force of 0.05 N (lower than the minimum 0.2 N used in Experiment 1), the typical 0.4 N (as in Experiment 1), and a higher force of 1.5 N (lower than the maximum 2.0 N used in Experiment 1). This was carried out for two reasons: the forces corresponded to those used in an earlier psychophysical study, in which some of the microneurography was conducted in conjunction with (Ali *et al*., 2023), and that fewer forces could be tested in microneurography due to experimental constraints (e.g. duration of unit recordings, the application of higher forces could pull too much on the skin and disrupt the recording). Therefore, the lower force of 0.05 N was used as a minimal level to explore whether C-LTMRs responded to such low stroking forces and the upper force allowed testing of a high force without compromising experimental stability. Further, instead of a soft brush, a flat-bottomed probe covered by a soft polyurethane foam (stroking surface ∼10 x 2 cm) was used, as per Ali *et al*., (2023). A comparison of the firing of C-LTMRs in previous work between a soft brush and a smooth metal plate at skin temperature found a complete overlap of C-LTMR responses to these stroking velocities, therefore we expected equivalence using a soft foam surface (Löken *et al*., 2009; Ackerley, Wasling *et al*., 2014; Watkins, 2022). The stimuli were applied in a pseudorandomised order and each velocity-force combination gave a total of 15 applied stimuli.

#### Procedure

The set-up was similar to Experiment 1, where participants sat comfortably in a reclining chair with their left arm immobilised with a vacuum cushion. Perceptual ratings were not obtained during microneurography, as per previous experiments (Löken *et al*., 2009; Ackerley, Wasling *et al*., 2014), as the microneurography approach is already challenging and the aim for Experiment 2 was to obtain electrophysiological recordings only. Peripheral nerve differential axonal recordings were made via microneurography (Ackerley & Watkins, 2022), where an insulated high impedance electrode (FHC, Bowdoin) was inserted into either the lateral antebrachial nerve ∼3 cm proximal to the cubital fossa (n = 4 participants) or into the radial nerve at the upper arm (n = 1 participant) to record from the hairy skin of the lower arm. Neural signals were amplified, visualised, and recorded using a NeuroAmpX connected to a PowerAmp with LabChart software (ADInstruments, Australia). Single LTMR units were identified via skin caressing, where C-LTMRs were classified according to Ackerley (2022), which included having a low threshold to indentation from a calibrated monofilament and high sensitivity to gentle stroking with a soft brush. Once a single C-LTMR of sufficient signal-to-noise had been identified and characterised, the RTS was positioned perpendicularly over the middle of the receptive field, the chosen forces (0.05 – 1.5 N) were calibrated, and the protocol applied.

#### Analysis

Microneurography data were analysed using LabChart (ADInstruments, Australia) and SpikeSort implemented in MATLAB (University of Gothenburg, Sweden). Single spike impulses were identified via a template-matching procedure and time-stamped. Data analysis on the time-stamped spikes was performed using RStudio, as per the ratings and RTS force data. In brief, for the analysis of the microneurography C-LTMR mean frequencies, a generalized linear mixed model was used due to the non-normal distribution of the data, where firing frequency was set as the dependant variable, stroking velocity and set force as fixed factors with their interaction, and each C-LTMR unit as a random factor, including a random intercept per unit. Bonferroni corrected post-hoc tests were conducted per force and per velocity.

## Results

### Experiment 1: Psychophysical ratings to different stroking velocities and forces

#### Perceptual pleasantness ratings

We first explored whether the relationship between tactile pleasantness and stroking velocity was best fit by linear or quadratic regressions at each force. For all forces, there was a significant negative quadratic fit (p < 0.001 at 0.2, 0.4, 0.8, 1.2 and 1.6 N, and p = 0.0226 at 2N) indicating an inverted-U shaped curve, but for the lower force curves from 0.2 to 1.2 N, a positive linear fit was also significant (p = 0.00553 at 0.2 N, p = 0.00241 at 0.4 N, p < 0.001 at 0.8 N, p = 0.0310 at 1.2 N, while p = 0.413 at 1.6 N and p = 0.278 at 2 N). This can be seen in Figure 2, where pleasantness increases for all forces during stroking at 0.3 to 1 cm.s^−1^. There is an overall peak in pleasantness around 3-10 cm.s^−1^ in the mean values over stroking velocities shown in Fig. 2B, yet this is clearly dependent on force, where forces over 1.2 N were much less pleasant, as seen in the pleasantness means over forces in Fig. 2C.

**Figure 2:**
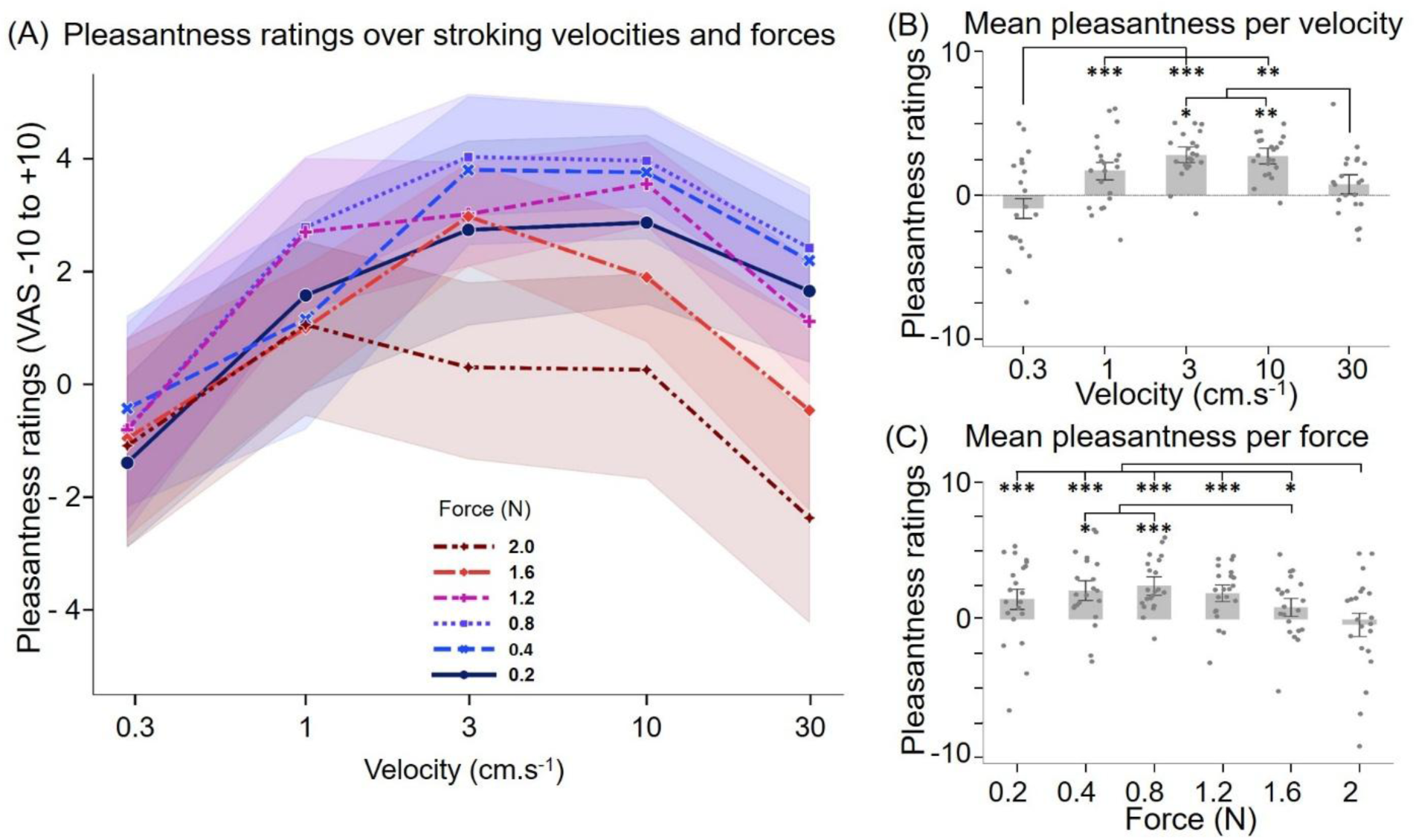
Pleasantness ratings are influenced by the stroking velocity and applied force. (A) Pleasantness ratings on a visual analogue scale (VAS) from −10 (very unpleasant) to +10 (very pleasant) over stroking velocities (x-axis) and forces (different lines), showing the means with ±95% confidence intervals. (B) Pleasantness ratings over velocities, averaged by force. Means with ±95% confidence intervals and individual participant ratings are shown. Significant differences between velocities were found at 0.3 cm.s^−1^, which was significantly less pleasant than all the other velocities, and at 30 cm.s^−1^, which was significantly lower than at 3 and 10 cm.s^−1^. (C) Pleasantness ratings over forces, averaged by velocity. Means with ±95% confidence intervals and individual participant ratings are shown. Significant decreases in pleasantness between 2.0 N and all other forces, apart from 1.6 N, were found, as well as a significant decrease in pleasantness was also found from 1.2 to 1.6 N. From n = 21 participants, *p < 0.05, **p < 0.01, ***p < 0.001.

To investigate differences between the stroking velocities and forces, we applied a linear mixed model that revealed a significant effect of stroking velocity (F_4, 34.87_ = 10.95, p < 0.001, η²ₚ = 0.56; Fig. 2B) and force (F_5, 540.0_ = 15.167, p < 0.001, η²ₚ = 0.12; Fig. 2C), as well as a significant interaction between these (F_20, 540.0_ = 1.89, p = 0.0107, η²ₚ = 0.07; Fig. 2A). For the effect of stroking velocity, Bonferroni-corrected post-hoc comparison tests showed that pleasantness at 0.3 cm.s^−1^ was significantly lower than all the other velocities (p < 0.001 compared to 1 and 3cm.s^−1^, p = 0.00452 compared to 10 cm.s^−1^), except 30 cm.s^−1^ (p = 0.715) and pleasantness at 30 cm.s^−1^ was significantly lower than 10 (p = 0.00303) and 3 cm.s^−1^ (p = 0.0107); Fig. 2B). For Force, mean pleasantness ratings were significantly higher for stroking at 2 N compared to 0.2, 0.4, 0.8 and 1.2 N (all p < 0.001) and to 1.6 N (p = 0.0124). 1.6 N was also significantly different from 0.4 (p = 0.0204) and 0.8 N (p < 0.001; Fig. 2C).

To investigate differences in the interaction between conditions, we conducted Bonferroni-corrected post-hoc contrast tests that revealed significant decreases in pleasantness for higher stroking velocities at higher forces. Table 1 summarises these differences, where this effect starts at 3 cm.s^−1^ and becomes greater the faster the velocity. At 10 and 30 cm.s^−1^ stroking, there are numerous significant decreases for higher force stroking (1.6 and 2.0 N), mainly showing differences between the intermediate forces of 0.8 and 1.2 N. Although no significant differences were found between the lowest forces and the intermediate forces, Figure 2 and Table 1 indicate that the intermediate forces were the most pleasant.

**Table 1:** Significant differences in pleasantness ratings over forces per stroking velocity. Pairwise contrasts per stroking velocity for pleasantness ratings, showing more significant differences between forces as velocity increased.

| **Velocity** | **Significant differences in force comparisons per velocity** |
| --- | --- |
| 0.3 cm.s<sup>-1</sup> | No significant differences between pleasantness ratings over forces: all p values = 1.000. |
| 1 cm.s<sup>-1</sup> | No significant differences between pleasantness ratings over forces: all p = 1.000, except for 0.4 N compared to 0.8 (p = 0.778) and 1.2 N (p = 0.982), for 1.6 N compared to 0.8 (p = 0.498) and 1.2 N (p = 0.638), and for 2 N compared to 0.8 (p = 0.589) and 1.2 N (p = 0.751). |
| 3 cm.s<sup>-1</sup> | 2.0 N less pleasant than 0.4, 0.8 (both p < 0.001), 1.2 (p = 0.0187) and 1.6 N (p = 0.0221). All other p values = 1.000, except for 2 N compared to 0.2 N (p = 0.0562). |
| 10 cm.s<sup>-1</sup> | 2.0 N less pleasant than 0.2 (p = 0.0285), 0.4 and 0.8 (both p < 0.001) as well as 1.2 N (p = 0.00141). All other p = 1.000, except for 2 N compared to 1.6 N (p = 0.752), and for 1.6 N compared to 0.4 (p = 0.402), 0.8 (p = 0.210), and 1.2 (p = 0.733). |
| 30 cm.s<sup>-1</sup> | 1.6 N less pleasant than 0.4 N (p = 0.0234) and 0.8 N (p = 0.00946)<br>2.0 N less pleasant than 0.4 N to 1.2 N (all p < 0.001). All other p values = 1.000 except for 2 N compared to 1.6 N (p = 0.347), 1.6 N compared to 0.2 (p = 0.176) and 1.2 N (p = 0.900). |

#### Perceptual intensity ratings

We next investigated tactile intensity perception and explored whether the velocity curves over the forces were best fit by linear or quadratic regressions. For all the forces, significant positive linear regressions fit the data (all p < 0.001), as can be clearly seen in Figure 3A where there was a monotonic increase in tactile intensity with stroking velocity and with force.

**Figure 3:**
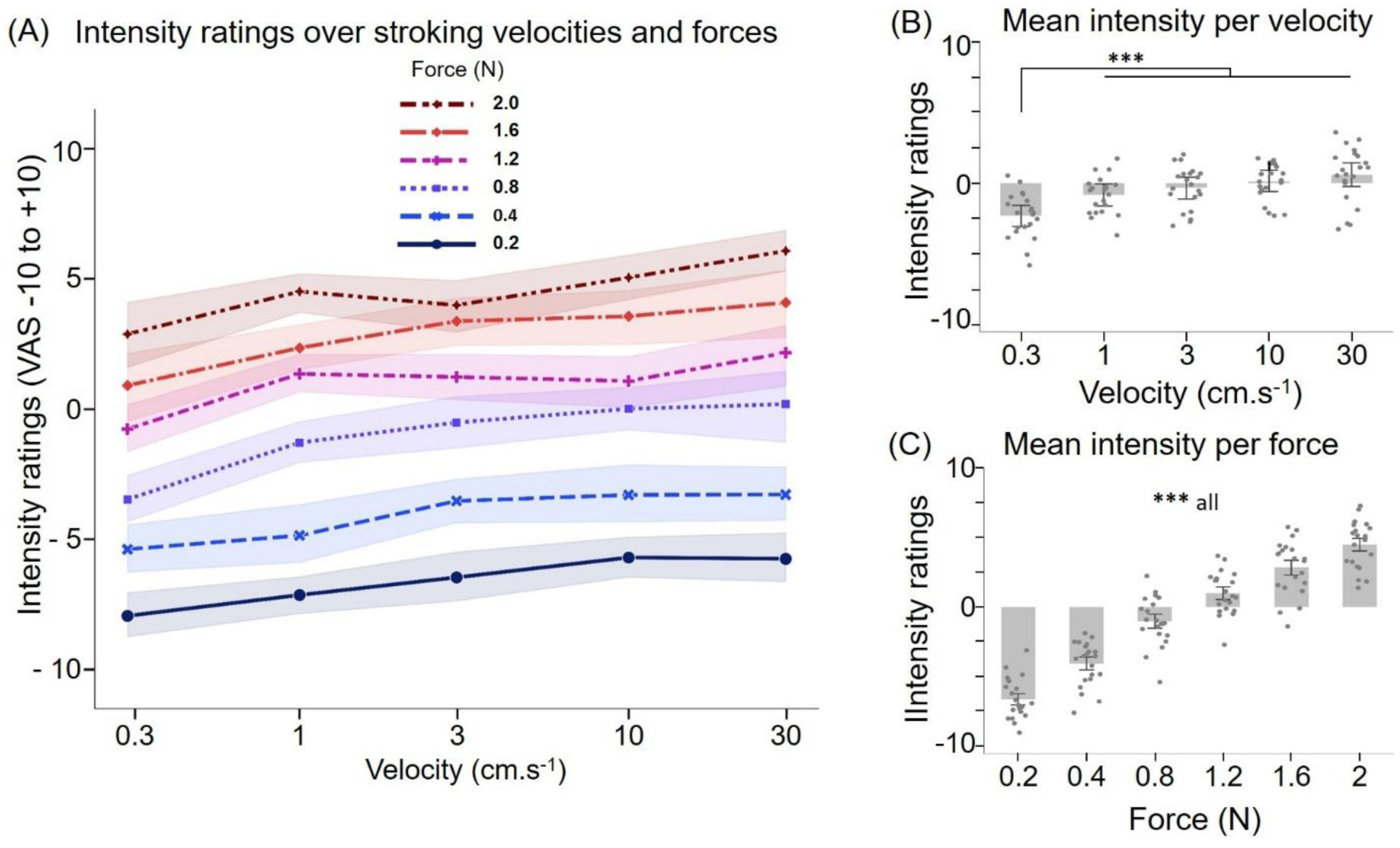
Intensity ratings are influenced by the stroking velocity and applied force. (A) Intensity ratings on a visual analogue scale (VAS) from −10 (very weak) to +10 (very strong) over stroking velocities (x-axis) and forces (different lines), showing the means with ±95% confidence intervals. (B) Intensity ratings over velocities, averaged by force. Means with ±95% confidence intervals and individual participant ratings are shown. Significant differences between velocities were found at 0.3 cm.s^−1^, which was significantly less intense than all the other velocities. (C) Intensity ratings over forces, averaged by velocity. Means with ±95% confidence intervals and individual participant ratings are shown. All comparisons were significantly different. From n = 21 participants, ***p < 0.001.

To investigate differences between the stroking velocities and forces, we used a linear mixed model that showed a significant effect of stroking velocity (F_4, 36.65_ =10.75, p < 0.001, η²ₚ = 0.54; Fig. 3B) and force (F_5, 539.45_ = 534.77, p < 0.001, η²ₚ = 0.83; Fig. 3C), with no significant interaction between these (F_20, 539.45_ = 1.07, p = 0.381; Fig. 3A). In Figure 3, which illustrates these effects, it is evident that participants were able to distinguish between all velocities and forces with respect to tactile intensity. For mean intensity over each velocity, Bonferroni-corrected post-hoc tests showed that 0.3 cm.s^−1^ was significantly lower than all other velocities (all p < 0.001; Fig. 3B). For force, all comparisons between all force levels were significantly different (all p < 0.001; Fig. 3C).

#### Comparisons of actual peak normal forces delivered over different stroking velocities and forces

We analysed the recorded force output curves from the RTS during each stroking trial to explore whether the forces delivered matched the set forces, as this is rarely shown (cf. Edin *et al*., 1995; Essick *et al*., 2010), but is important to consider in the present work. We found that the RTS delivered a systematic under-estimation of the set force, which became larger at the higher forces (Fig. 4). Figure 4A shows this under-estimation per force and Figure 4B illustrates each force level. There was a significant positive linear relationship for all the forces over the velocities (all p<0.001 except for 0.4 N where p = 0.00503), apart from no relationship at 0.2 N (p = 0.271). A linear mixed model gave significant main effects of velocity (F_4, 580_ = 106.14, p < 0.001, η²ₚ = 0.42) and set force (F_5,580_ = 8465.98, p < 0.001, η²ₚ = 0.99) were extremely strong, and there was a significant interaction between these (F_20, 580_ = 15.5, p < 0.001, η²ₚ = 0.35). Bonferroni-corrected post-hoc tests showed that all set forces were significantly different (all p<0.001) and all velocities were significantly different (all p<0.001 except for 1 compared to 0.3 cm.s^−1^ where p = 0.0352). To determine if there were differences for each force level over increasing stroking velocity, we compared RTS output forces at 30 cm.s^−1^ to the other velocities, for each set force. No significant differences were found between recorded normal forces applied at the 0.2 (all p values = 1.000) and 0.4 N (p = 1.000 compared to 10 cm.s^−1^, p = 0.408 compared to 3cm.s^−1^, p = 0.794 compared to 1 cm.s^−1^, p = 0.202 compared to 0.3 cm.s^−1^) set forces. At the set force of 0.8 N the actual force delivered was significantly higher at 30 cm.s^−1^ compared to 0.3 (p = 0.00161) and 1 cm^−1^ (p = 0.00668), but not compared to 10 (p = 1.000) and 3 cm.s^−1^ (p = 0.336). At the set force of 1.2 N, the actual force delivered was significantly higher at 30 cm.s^−1^ as compared to all other velocities (p < 0.001 compared to 0.3 and 1 cm.s^−1^, and p = 0.00141 compared to 3 cm.s^−1^), apart from 10 cm.s^− 1^ (where p = 0.677). At the highest set forces of 1.6 and 2.0 N, this effect was stronger, where there was a significant increase in delivered force at 30 cm.s^−1^, as compared to all the other velocities (all p < 0.001).

**Figure 4:**
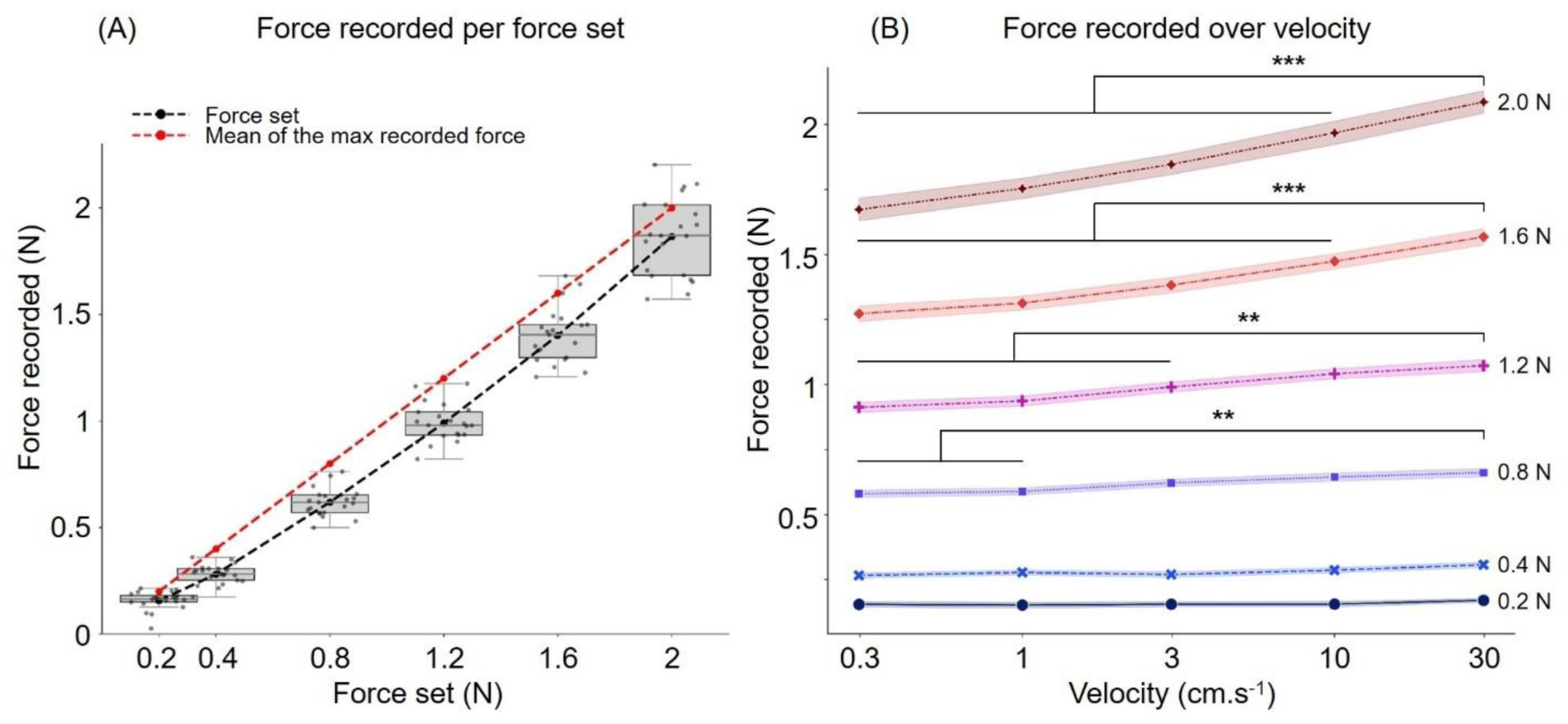
Comparisons of actual maximum normal forces delivered by the RTS. In the psychophysical study, we recorded and analysed the actual delivered maximum normal forces from the rotary tactile stimulator (RTS). (A) This shows the set forces (in red) and the actual peak normal forces obtained (in black, with box-and-whisker plots; n = 21 participants). (B) Actual peak normal forces over the different velocities per set force, showing means with ±95% confidence intervals.

### Experiment 2: Microneurography recordings of C-LTMR afferent responses to different stroking velocities and forces

Due to the challenges of microneurography, a total of seven C-LTMRs were obtained for the full stroking paradigm. We found that the C-LTMRs mean firing frequency increased with increasing stroking velocity and force (Fig. 5). The C-LTMRs had mechanical indentation thresholds of <0.005 N and their locations are shown in Figure 5B. Over the stroking velocities at 1.5 N, both significant positive linear (p = 0.0104) and negative quadratic (p = 0.0215) relationships were found, but no significant linear nor quadratic relationship at 0.05 (linear: p = 0.298, quadratic: p = 0.490), nor at 0.4 N (linear: p = 0.0539, quadratic: p = 0.898). A generalized linear mixed model gave a significant main effect of stroking velocity (χ² (4) = 10.29, p = 0.036) and force (χ² (2) = 15.76, p < 0.001), but no significant interaction between these (χ² (8) = 2.67, p = 0.953). Bonferroni-corrected post-hoc tests showed that the velocity 0.3 cm.s^−1^ was significantly different to 10 and 30 cm.s^−1^ (p = 0.00970 and p = 0.00562, respectively) and that all force levels were significantly different (all p < 0.001, apart from 0.4 N compared to 1.5 N, which was p = 0.0115).

**Figure 5:**
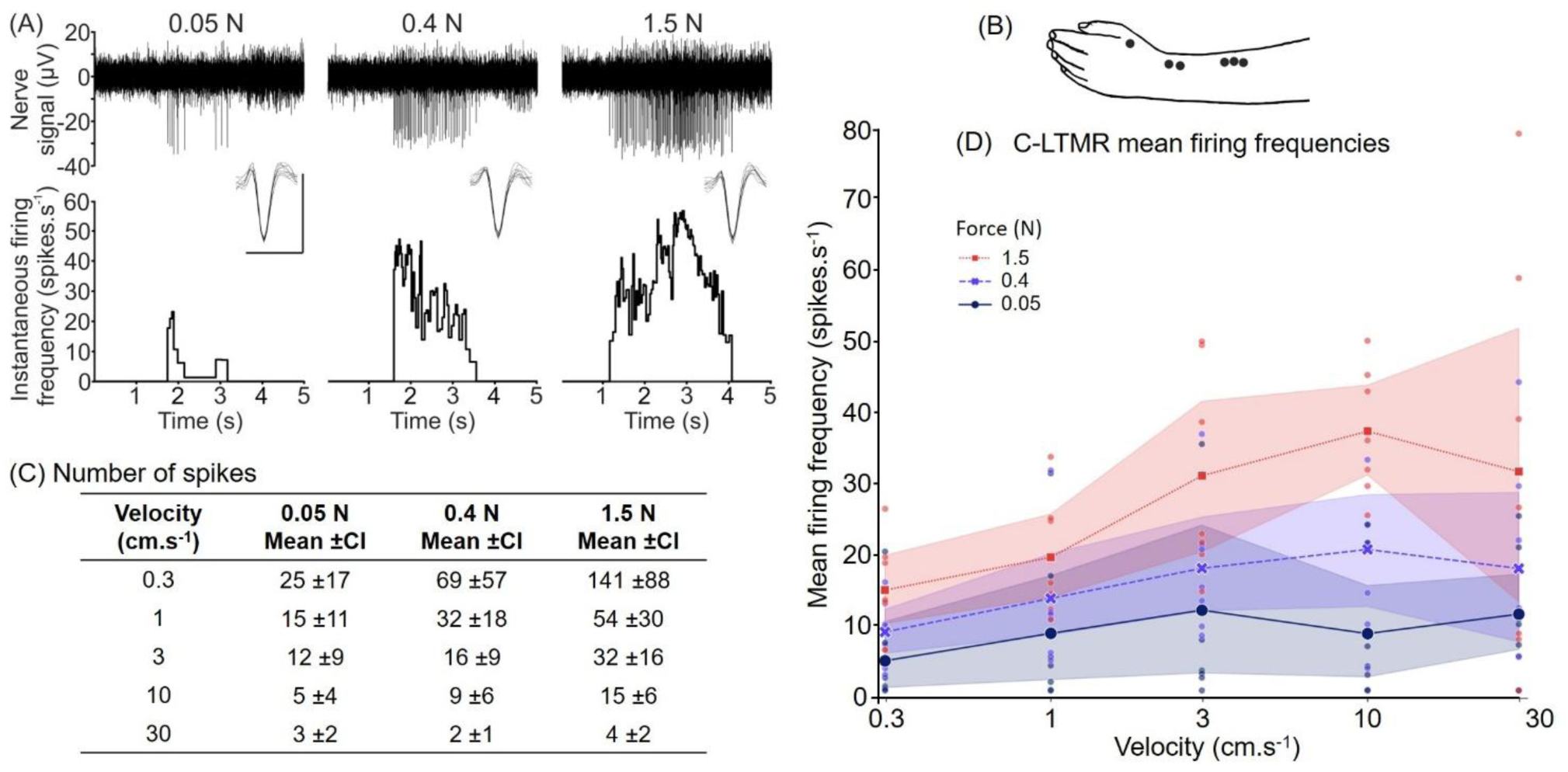
Firing frequencies of C-LTMRs over different stroking velocities and forces. (A) Examples of C-LTMR firing and the instantaneous firing frequency for one unit stroked at 1 cm.s^−1^ over 0.05, 0.4, and 1.5 N. The inset spikes show an overlay of nine spikes from each trace, with the scale being 50µV amplitude and 1 ms time. (B) Locations of the C-LTMR units. (C) Number of spikes found for each stroking velocity and force, giving the mean and 95% confidence intervals. (D) Mean firing frequency per stroking velocity at the forces of 0.05, 0.4, and 1.5 N (indicted by different lines and colours), with 95% confidence intervals as shading and smaller dots showing individual data points from n = 7 C-LTMRs.

## Discussion

In the present study, we investigated the effects of stroking velocity and force on the perception of touch and relate this to activity in C-LTMRs. Previous work has underscored the importance of stroking velocity in the perception of pleasantness, which has shown direct links to firing in C-LTMRs (Löken *et al*., 2009; Ackerley, Wasling *et al*., 2014), and we take this further by showing that force is an important factor in both tactile pleasantness and intensity evaluations, as well as having a strong effect on C-LTMR activity. Both stroking velocity and force significantly influenced pleasantness and intensity ratings, although different effects were found for each. For pleasantness, the typical inverted-U shaped relationship was found for stroking velocities (Croy *et al*., 2021) and although a similar relationship was found for force, these two factors interacted to give significantly lower pleasantness when the stroke was fast and forceful. No effect of force was found during very slow stroking, but the effect became more pronounced the faster the stroke became. Intensity ratings followed clear linear relationships, where both increases in stroking velocity and force produced increased tactile intensity. The recordings from C-LTMRs highlight that neither the pleasantness, nor intensity, perceptual results mapped directly onto the activity of C-LTMRs, and we discuss the implications below.

Regarding the pleasantness findings, our results confirm the numerous studies that have shown an inverted-U shape between pleasantness and stroking velocity, with very slow and very fast velocities being slightly unpleasant, and a peak in pleasantness between 1-10 cm.s^−1^ (Essick *et al*., 1999; Löken *et al*., 2009; Ackerley, Carlsson *et al*., 2014; Ackerley, Wasling *et al*., 2014; Croy *et al*., 2021). This is a relatively robust finding, although there is inherent variability in the ratings and this relationship is not always found at an individual level, where many participants show positive linear curves as well (Croy *et al*., 2021). Stroking force has been little investigated in pleasantness perception, where the majority of studies have used a force of ∼0.4 N, although this is only well-controlled using a stroking robot. Löken *et al*. (2009) tested stroking forces of 0.2 and 0.4 N, but found no significant differences in terms of pleasantness, nor C-LTMR firing; however, the forces did not differ great and only two were tested. Other studies have investigated different forces and textures, finding a general decrease in pleasantness with force, but it was less clear for smooth surfaces (Cascio *et al*., 2008; Essick *et al*., 2010; Ali *et al*., 2023). We extend these findings by investigating numerous stroking velocities and forces that were robot-applied to show that even with brush stroking, pleasantness decreases with force, but only at faster stroking velocities. This implies that stroking velocity and force give related pleasantness percepts, but that there is an interaction, potentially due to the relative encoding by mechanoreceptors.

To complement the pleasantness ratings, we also investigated tactile intensity. In general, the literature shows that tactile intensity increases linearly with stroking velocity (Triscoli *et al*., 2013; Jönsson *et al*., 2015; Sehlstedt *et al*., 2016; Schirmer, Cham *et al*., 2023), as we presently found. Akin pleasantness ratings, tactile intensity was also influenced by force, but increased linearly with it, where all comparisons between forces were significantly different. Similar results have been found previously between three forces, although no significant interaction was found between stroking velocity and force (Ali *et al*., 2023), which we currently found. The previous lack of interaction may have been due to fewer forces being tested, but in examining the present results touch seemed to be much less intense at both lower velocities and forces. Again, this may relate to the underlying encoding via mechanoreceptors.

The literature on Aβ-LTMRs shows their firing frequency increases exponentially with stroking velocity (Essick & Edin, 1995; Edin *et al*., 1995; Löken *et al*., 2009; Ackerley, Wasling *et al*., 2014), but this relationship does not map directly onto pleasantness (inverted-U shape), nor intensity (linear, with larger increases in the slope at lower velocities) ratings. This demonstrates how touch is a ‘team effort’ and the contributions of all types of sensory afferents can shape tactile perception (Saal & Bensmaia, 2014). However, it is nevertheless of interest to relate the firing in different afferent populations to tactile percepts and it can be seen that in general, tactile intensity does relate to increases in firing frequency in Aβ-LTMRs. When it comes to the contribution of C-LTMR afferents to tactile percepts, the relationships become more complex. C-LTMR input inherently arrives with a relative delay in the brain (seconds timescale), as compared to Aβ-LTMR input (milliseconds timescale), due to their slower conduction velocity. C-LTMRs have been postulated to play a direct role in signalling the affective and emotional aspects of touch, due to the relationship between their firing frequency and stroking velocity, which correlates with pleasantness ratings (Löken *et al*., 2009; Ackerley, Wasling *et al*., 2014). However, the firing of C-LTMRs does not always significantly correlate with perceived pleasantness when additional parameters are manipulated, such as stroking at temperatures that are cooler and warmer than skin temperature, where both the C-LTMR firing and pleasantness perception diverge (Ackerley, Wasling *et al*., 2014).

Therefore, although C-LTMR activity has been implicated in pleasantness perception, this is a complex process that also likely depends on other afferent signals, as well as central mechanisms (Vallbo *et al*., 2009; Morrison, 2022; Case *et al*., 2023; Schirmer, Croy *et al*., 2023). With regards to the encoding of force by C-LTMRs, few studies have investigated this, therefore we provide new insights into the encoding of moving touch by C-LTMRs that are increasingly activated by force. It is unsurprising that no significant differences were found between stroking at 0.2 and 0.4 N in Löken *et al*. (2009), as these are both relatively low forces. In more recent work, it was clearly shown that C-LTMRs encode statically-applied force, via calibrated monofilaments, from 0.004 to 0.1 N in a positive linear way (Middleton *et al*., 2022, supplementary figure 4B). However, the local forces produced from indentation are rather different to moving across the receptive field. Yet, we presently show that C-LTMRs encode different stroking forces well, possibly better than they encode stroking velocity.

Although we only obtained recordings from a total of 7 C-LTMRs for the full paradigm, the sample size is equivalent to other similar papers for LTMR recordings (Löken *et al*., 2009, 2022; Ackerley, Wasling *et al*., 2014), in part, due to the challenges of such microneurography experiments. Therefore, although further work should be conducted to explore dynamic force encoding in LTMRs more, the present results demonstrate clearly the capacity of C-LTMRs to signal force, although recordings from a wider range of forces would be ideal. Our results showed that C-LTMR firing increased to stroking velocities of 10 cm.s^−1^, but that the inverted-U shape curve was not evident. This does correspond somewhat to the previous work, where different mechanisms underlie the responses of C-LTMRs to each side of the stroking curve (Ackerley, 2022). At very slow velocities, C-LTMRs fire a lot, it is just that their mean firing frequency is overall lower than at intermediate velocities (Löken *et al*., 2009; Ackerley *et al*., 2018). Conversely, C-LTMRs reach their limit of velocity encoding at 30 cm.s^−1^, where there is either no response or a few spikes; only when at least two spikes are present can a firing frequency be determined, hence the high variability seen in our results at 30 cm.s^−1^. The inverted-U shape velocity tuning was not entirely present, although not unexpected and adding more C-LTMRs could help see this better, the effect of stroking force dominated. C-LTMRs clearly fired much more to 1.5 N, as compared to stroking at lower forces, yet we can ask whether pleasantness was directly related to this. Our perceptual results showed highest pleasantness for 0.2 to 0.8 N stroking, with no differences between these forces, therefore the activity tuning to force in C-LTMRs is intriguing. However, this again underlies the idea that touch depends on all incoming inputs, from A- and C-fibre mechanoreceptive afferents, to other afferents such as thermoreceptors and nociceptors. Further work should investigate the effects of forces more on different Aβ-LTMRs, as although Aβ-LTMRs have been shown to encode force monotonically, it is possible that various types encode forces in different ways. It is also likely that at higher forces, C-mechanosensitive nociceptors start to contribute to perception. Studies have shown that C-mechanosensitive nociceptors can occasionally be activated via monofilament indentation from 0.0025 N and can fire occasionally to a brush stroke (Watkins *et al*., 2017). Therefore, with increasing force, these may be recruited more and the balance is tipped from C-LTMR-dominating positive affect to increased nociceptor negative affect, which should be investigated in further work.

As well as the need to record from more afferents, further work could probe the effect that we found of increasing RTS applied force with increased stroking velocity. Overall, the recorded force was always less than the set force and we found that this effect was more profound at slower stroking and higher forces. Few studies have looked at the output from the RTS, which is calibrated in terms of stroking direction, speed, and force. However, the significant increase in delivered, as compared to set, normal force may have impacted the perception. It is difficult to quantify the exact impact in the present study, therefore future work should take this into consideration, both mechanically and perceptually, when exploring interactions over the skin.

To conclude, our results call for a revision of current models of affective touch perception in which force is often not considered a central parameter. This relates well to more recent work investigating the pleasantness of deep touch, which is an important aspect in affective and social touch, such as the impact of a hug or relaxation from a massage (Case *et al*., 2020, 2023). Both tactile pleasantness and intensity clearly relate to the firing of C-LTMR and Aβ-LTMR afferents, respectively, yet mounting evidence, combined with our present findings, demonstrate the complexity of affective touch perception and that it requires input from many different types of mechanoreceptors and other sensory afferents.

## Generative AI Statement

No generative AI tools were used in the preparation of this manuscript.

## Data Availability

Data and scripts for Experiment 1 are available via an OSF repository at: https://osf.io/gxnyj/overview?view_only=26bb90dda8e04b4f99a7f1b1fa128148. Due to the more sensitive nature of human microneurography recordings, the microneurography data can be accessed upon reasonable request.

## Author Contributions

S.F., L.T., and R.A. designed, performed, and analysed Experiment 1. For Experiment 2, P.D.T., A.G.M., A.M., S.H.A., and F.M. designed the study, A.M., A.G.M., and S.H.A. performed the study, A.M. and S.H.A. analysed the data, and P.D.T. supervised the study. S.F. drafted the first version of the manuscript with R.A. All authors approved the final version of the manuscript. All authors agree to be accountable for all aspects of the work in ensuring that questions related to the accuracy or integrity of any part of the work are appropriately investigated and resolved. All persons designated as authors qualify for authorship, and all those who qualify for authorship are listed.

## Acknowledgments

S.F. was funded by the CNRS MITI program in a grant to R.A and by the Fondation ARC. The study was funded by an Agence National de la Recherche (ANR, France) grant ‘REMASS’ to R.A. in the CRCNS International call on Computational Neuroscience – NEUC 2024 collaboration. A.G.M was funded by the Pain Relief Foundation. This work was supported by an Undergraduate Research and Knowledge Exchange Internship Scheme (URKEIS) award from Manchester Metropolitan University for which S.H.A. was the research intern.

